# A Workflow of Integrated Resources to Catalyze Network Pharmacology Driven COVID-19 Research

**DOI:** 10.1101/2020.11.04.369041

**Authors:** Gergely Zahoránszky-Kőhalmi, Vishal B. Siramshetty, Praveen Kumar, Manideep Gurumurthy, Busola Grillo, Biju Mathew, Dimitrios Metaxatos, Mark Backus, Tim Mierzwa, Reid Simon, Ivan Grishagin, Laura Brovold, Ewy A. Mathé, Matthew D. Hall, Samuel G. Michael, Alexander G. Godfrey, Jordi Mestres, Lars J. Jensen, Tudor I. Oprea

## Abstract

**Motivation:** In the event of an outbreak due to an emerging pathogen, time is of the essence to contain or to mitigate the spread of the disease. Drug repositioning is one of the strategies that has the potential to deliver therapeutics relatively quickly. The SARS-CoV-2 pandemic has shown that integrating critical data resources to drive drug-repositioning studies, involving host-host, hostpathogen and drug-target interactions, remains a time-consuming effort that translates to a delay in the development and delivery of a life-saving therapy.

**Results:** Here, we describe a workflow we designed for a semi-automated integration of rapidly emerging datasets that can be generally adopted in a broad network pharmacology research setting. The workflow was used to construct a COVID-19 focused multimodal network that integrates 487 host-pathogen, 74,805 host-host protein and 1,265 drug-target interactions. The resultant Neo4j graph database named “Neo4COVID19” is accessible via a web interface and via API calls based on the Bolt protocol. We believe that our Neo4COVID19 database will be a valuable asset to the research community and will catalyze the discovery of therapeutics to fight COVID-19.

**Availability:** https://neo4covid19.ncats.io

## Introduction

The pandemic of the SARS-CoV-2 virus (also commonly referred to as COVID-19 pandemic by the name of the disease it induces) put a spotlight on the need for mechanisms that can rapidly identify and integrate relevant information to give a fighting chance to the medical and research community. The Ebola and Zika outbreaks presented similar challenges, and significant advances have been made in the past years in terms of digital, laboratory, and epidemiological techniques [1]–[3]. Although numerous resources covering many aspects of pertinent biomedical research have been developed and made publicly available, e.g. ChEMBL[4], [5], Reactome [6], Pharos [7], PathwayCommons [8], BioPlanet [9], and DrugCentral [10] to name a few, their on-demand integration has been a translational bottleneck to date.

In the case of the outbreak of an unknown pathogen, such as SARS-CoV-2, the first line of defense – absent viable therapeutic options – is containment. Should this fail, we have to resort to mitigation to slow down the spread of the pathogen. Delays of mere days in the early stages of containment and mitigation can lead to catastrophic outcomes regarding the number of infections and death toll [11]–[13]. Furthermore, the COVID-19 pandemic was reported to have a dramatic impact on primary healthcare providers, limiting them to essential clinical services, which eventually led to unforeseen delays in diagnosing highly critical diseases such as cancers [14], [15]. Therefore, it is imperative that we have computational workflows in place that can help researchers to connect and navigate the relevant information *very* fast – *much* faster than today. Multiple pertinent datasets subjected to such a workflow would produce a condensed, enriched starting point for timely hypothesis generation that would drive a successful containment or mitigation strategy, such as drug repositioning [16].

Indeed, a small number of publicly available databases and knowledge-graphs have been reported that connect various types of information related to COVID-19. Such resources are primarily limited to data extracted via text mining from literature and patents, reports, and experimental data [17]–[26]. While these resources are valuable for advancing our efforts toward a possible therapy for COVID-19, they suffer from certain limitations from a translational point of view.

First, hypothesis generation in the current drug discovery paradigm, i.e. network pharmacology [27]–[29], can be enhanced by normalizing the nature of interactions between protein target pairs, and between compounds and targets to reflect if they are engaged in a stimulatory or an inhibitory relationship. Although necessary information might be present in existing data sources, the interaction categories are typically encoded as separate relationships. This makes it difficult to readily assess the inhibitory and stimulatory relationships. For instance, both “antagonist” and “ion channel blocker” actions can be further reduced to an “inhibitory” relationship to aid the analysis of network perturbation. Naturally, the original relationships can also be preserved to avoid loss of information. A normalized relationship of protein-protein and drug-target interactions would consist of only three values: “stimulates”, “inhibits”, and “undefined” (or the equivalent of these phrases).

Next, the recent concept of “target development level (TDL)” [7] of protein targets is not captured in existing knowledge graphs related to COVID-19. The annotation of TDL category of targets makes it easy to identify those whose activity can be modulated by FDA-approved drugs or by small molecules. Such information therefore facilitates drug repositioning-oriented hypothesis generation.

Another category of limitations pertains to translational aspects: knowledge dissemination and real-time data integration. To our knowledge, no data source related to COVID-19 to date has been equipped with a mechanism to facilitate the data exploration and analysis for those without and with substantial bioinformatics background at the same time. However, in the case of a pandemic, it is imperative to disseminate data sources and data analysis tools to as broad a scientific community as possible and as soon as possible.

Finally, from a technical standpoint, it is of paramount importance that we have publicly available mechanisms and workflows for real-time integration of heterogeneous information. Typically, careful integration of databases can take months and even years, and oftentimes the integration workflow is quite specific for the knowledge base at hand [30], [31].

In this study, we describe such a workflow and utilize it to assemble, from several pertinent sources, a knowledge network aimed at defeating COVID-19 via enhanced hypothesis generation. The first building block is a list of pathogen–host protein interactions that was published in a preprint on March 27, 2020 by Krogan *et al.* [32], [33] within two months of the disclosure of the SARS-CoV-2 genetic sequence and within two weeks of the WHO declaring COVID-19 a pandemic [34], [35]. Further building blocks encompass interactions between FDA approved drugs and host protein targets (DTIs), host-host and host–pathogen protein-protein interactions (HHIs, and HPIs, respectively), and predicted drugs and host targets that will be introduced in this study. While this particular resultant Neo4j database is intended to catalyze COVID-19 research, the process of its creation can serve as a blueprint for inevitable future outbreaks, translating to saving precious time and, consequently, lives.

## Related Work

The workflow presented in this study was inspired by earlier works, namely SmartGraph [36] and Hetionet [37], both of which are so-called multimodal networks designed to aid drug discovery efforts in a network pharmacology setting. SmartGraph is a computational platform that consist of a knowledge base and a web-based user interface. The knowledge base of SmartGraph integrates drug-target and protein-protein interactions. The user interface integrates complex but easy-to-execute workflows for the analysis of network perturbation, and for bioactivity prediction and drug repositioning. Relationships between proteins are labeled as stimulatory or inhibitory to reflect the nature of the interactions. Hetionet is an open-source resource that is of notable importance and relevance. A wide variety of publicly available biomedical, disease-specific, and pharmacological databases were integrated into a large network. Unlike SmartGraph, Hetionet annotates drug-target interactions in terms of pharmacological action. Nevertheless, neither SmartGraph nor Hetionet provide a normalized annotation of the stimulatory and inhibitory relationships between proteins or between drugs and targets.

In a recent effort, researchers from University of Minnesota, Hunan University, and Amazon’s AWS artificial intelligence laboratories have collectively built the COVID-19-related Drug Repurposing Knowledge Graph (DRKG) [25] that integrates Hetionet among other data sources. Although DRKG addresses, to some extent, the issue of knowledge dissemination, this particular resource is available only as data tables, a format that requires advanced bioinformatics skills for analyzing the data in terms of network perturbation caused by drugs. Furthermore, the workflow to assemble the DRKG is not publicly available. COVID-KG [24] is another resource that focuses on extracting multimedia knowledge elements from scientific literature to be used as a knowledge graph for querying and report generation. While researchers developing both DRKG and COVID-KG have addressed the issue of data dissemination to some extent, the format they chose for distribution (tab-separated flat files) restricts network-based data analytics absent advanced bioinformatics skills. Furthermore, neither DRKG nor COVID-KG are associated with a publicly available implementation reflective of their underlying data integration workflows.

Another COVID-19 related resource is the recent “COVID-19 Disease Map” that was created by the means of a collaborative effort. While this study addresses the importance of the standardization of file formats, it does not provide a flexible workflow (and implementation as source code) that could serve as the basis for an automated or semi-automated data integration process. Although at the time writing of this manuscript the COVID-19 Disease Map is not yet accessible, the description of the integration process indicates that much emphasis was put on the use of curated data sources. This study exemplifies that a collaborative data integration strategy can yield high quality data sources, however, at the sacrifice of time.

Apart from KGs and databases, several other studies identified potential targets and contributed useful datasets that can guide drug discovery efforts. For instance, Gil *et al.* [17] provided their perspective on the main targets that are involved in viral replication and control of host cellular processes. In parallel, structure-based efforts [19], [20] have been reported where the authors performed molecular docking simulations and virtual high-throughput screening to prioritize candidates for drug repurposing. Experimental high-throughput screening data have been made available by the National Center for Advancing Translational Sciences (NCATS/NIH) on the “OpenData Portal” [21].

## Computational Methods and Datasets

### Assembly of a Multimodal Network

In a drug discovery setting driven by network pharmacology, a multimodal network [36]–[38] needs to be assembled and tailored according to the disease context. In this study, we set forth to create a multimodal network focused on COVID-19 by integrating host and pathogen protein targets, drugs, and relations defined between them, such as HHIs, HPIs and DTIs. The respective information was derived from various data sources described in detail below. Furthermore, we use a “data track” alias (see: *Fig. 1.*) for each dataset throughout the text in order to concisely and unambiguously refer to individual input datasets. A pseudo-code of the data integration workflow is provided in *“Pseudo-Code of the Data Integration Workflow”* section in Supporting Information (SI).

**Figure 1.**
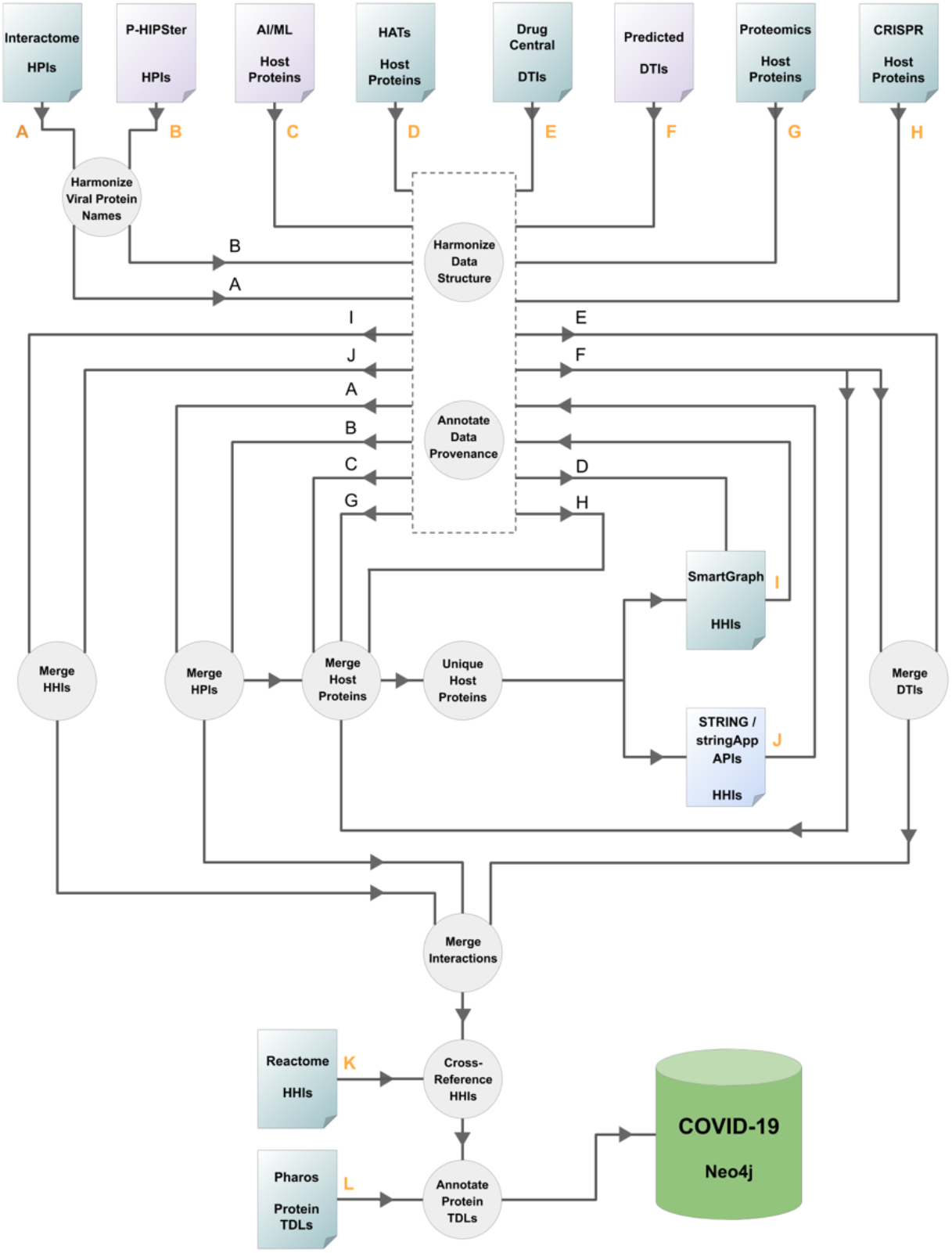
Resource integration logic. The schema highlights the most important steps of data processing. Individual inputs are labeled with letters. Orange letters indicate points of the workflow where new data sources are introduced. Arrowheads and letters aid to track the flow of information when it is not obvious. Purple, green and blue colors distinguish types of resources that utilize experimental, predicted and both types of data, respectively. HHIs: host–host protein interactions, HPIs: host–pathogen (here: SARS-CoV-2) protein interactions, DTIs: drug–target interactions, TDLs: target development levels.

### Host Targets Implicated to Play Important Role in Pathogenesis

#### Proteomics Study of Infected Cells

Host cell translation changes [39] following SARS-CoV-2 infection were studied in Caco-2 cells using translatome [40] and proteome proteomics at four different time points (2, 6, 10, and 24 hours, respectively) after infection. This unbiased profile of the cellular response to SARS-CoV-2 infection was used to identify key determinants of the host cell response to infection. Extensive proteome modulation occurs 24 hours post infection, e.g., reduced expression of cholesterol metabolism proteins, and increased expression profile for carbon metabolism proteins and spliceosome components. Some pathways appear amenable to therapeutic intervention, e.g., along the proteostasis and nucleotide biosynthesis pathways. Given quantified translation data for 2,715 proteins (as documented in the Supplementary files), we selected proteins with P values below 0.05 at 24 hours (virus exposure compared to control), as follows: 75 proteins having lower translation values across all 4 time points (of these, 38 are involved in acetylation according to STRING); and 23 proteins having positive virus-induced translation values at 24 hours (of which, 12 are also involved in acetylation), respectively. These 98 host proteins were subject to further processing: one (UniProt AC: Q9N2J8) was removed as it cannot be mapped to a gene name, four (UniProt ACs: P84243, Q8IZP9, Q8IXH7, P63302) were mapped to multiples gene names, thus the respective records were replicated. This gave rise to 102 proteins in total, that were denoted as data track “G”.

#### Genome-Wide CRISPR Study

Host genes essential for cell survival in response to SARS-CoV-2 infection were identified using two Cas9 Vero-E6 cell line constructs, using a genome-wide pooled CRISPR (clustered regularly interspaced short palindromic repeats) library [41], [42]. This CRISPR-Cas9 screen identified “pro-viral” and “anti-viral” genes, as follows. “Pro-viral” genes are involved in resistance, and their knockout confers resistance to virus-induced cell death. These genes include the ACE2 viral entry receptor, 11 genes from the SWI/SNF (SWItch/Sucrose Non-Fermentable) chromatin remodeling complex [43] and 7 genes associated with CDKN1A transcription upregulation via RUNX3 [44], respectively. “Anti-viral” genes are involved in sensitization, and their knockout sensitizes a cell to virus-induced cell death; these genes include HIRA (a subunit of the H3.3 histone chaperone complex), a set of 6 genes involved in viral translation, 8 genes associated with the SMN (survival of motor neurons) complex, and 5 components of the NURF (Nucleosome Remodeling Factor) complex, respectively. A set of 53 “pro-viral” and 52 “anti-viral” host genes detected via this CRISPR-Cas9 screen [42] were incorporated, and assigned a data track label “H”.

#### PredictedHost Proteins

Here we describe an AI/ML framework that was utilized to predict host proteins that potentially interact with viral proteins.

The Target Central Relational Database (TCRD) [7] aggregates protein-specific data from different sources (e.g., GTEx [45], LINCS [46], STRING [47], Reactome [6], [48]), and it was used to build the TCRD-KG knowledge graph. The TCRD-KG nodes can be proteins, diseases, or phenotypes, and edges can be pathways, protein-protein interactions, or other biological relationships among proteins and diseases.

A Machine Learning (ML) framework based on the TCRD-KG meta paths [49] and XGBoost [50] classification algorithm was developed to predict disease-associated genes (proteins). The meta paths specify network paths that connect proteins to specific diseases in the TCRD-KG. The degree-weighted path count (DWPC) [51] metric is used to quantify the meta paths prevalence, transforming TCRD-KG data to feature vectors for ML model by meta paths matching, based on an input disease. The Python package to build TCRD-KG using TCRD database and XGBoost ML model is available on GitHub (https://github.com/unmtransinfo/ProteinGraphML) [52].

A training data set comprising 104 proteins as positive labels (known to be associated with SARS-CoV-2) and 114 proteins as negative labels (not associated with SARS-CoV-2) was used to train the ML model. A total of six different models were built, using slight variations of the input data (e.g., inclusion/deletion of human proteins identified by P-HIPSTER [53] to interact with SARS-CoV-2 proteins; and inclusion or absence of the LINCS [46] descriptors). Using 5-fold cross-validation on the training data, the area under the curve (AUC), accuracy, and Matthews Correlation Coefficient (MCC) were computed for each model. The ML models trained on 218 proteins were used to predict the association between 20,029 proteins and SARS-CoV-2. Using XGBoost feature importance output, meta paths were sorted in decreasing order of “gain” score to understand node interactions. The features with higher “gain” were more dominant in predicting SARS-CoV-2 associated proteins. A total of 986 proteins were predicted with “high confidence” by the 6 models; of the 136 predicted by 3 or more models, 99 were not part of the “understudied” proteins [54] and were given preference for this study (denoted as data track “C”).

### Virus-Implicated Host Proteins

Host proteins that were either identified as interacting partners of SARS-CoV-2 virus proteins in experimental studies or were predicted to be of potential importance in terms of pathogenesis or therapy are referred to as virus-implicated host proteins (VIHPs) in this study. VIHPs were derived from experimental determined HPIs (data track “A”), and from host proteins implicated in experimental studies (data track “G”, “H”). This initial set of VIHPs was extended by predicted HPIs (data track “B”), DTIs (data track “F”) and host proteins of potential importance (“C”).

### Host–Pathogen and Host–Host Interactions

#### Experimentally Determined HPIs

Experimentally determined host–pathogen interactome was published by Krogan *et al.* [32], [33] They identified 332 HPIs defined between 26 viral and 332 host proteins or host factors (data track “A”) that were determined via affinity purification mass-spectroscopy (AP-MS) [54]. A systematic analysis revealed 67 druggable human proteins and 69 compounds (including FDA-approved drugs and investigational drugs currently tested in clinical trials and preclinical studies) that were proposed to be evaluated for efficacy against SARS-CoV-2. While the host–pathogen interactome suggests potential pharmacological targets and possible interventions, the outcomes should be cautiously interpreted as it was mentioned that the identified agents could have either beneficial or detrimental effects (e.g. HDAC2 inhibitors).

#### InferredHPIs

Potential HPIs were predicted using the P-HIPSTer algorithm [53] giving rise to 155 HPIs involving 28 SARS-CoV-2 and 38 human proteins (data track “B”). P-HIPSTer offers a computational framework that utilizes sequence and structural information to infer HPIs, currently covering a total of 28 viral families (>900 human viruses) and > 5,000 human proteins accounting for 282,528 HPIs [55].

#### Experimentally Determined HHIs

We collected HHIs from various data sources. The network assembly process involved a network expansion step with the help of the STRING and stringApp APIs (data track “J”) [47], [56]. In this step, experimentally determined and inferred HHIs were imported with the following parameter settings: maximum interactors = 100, alpha = 0.5. For more details on STRING expansion of the network please refer to section *“Expansion of HHIs via StringApp API”* in SI. Considering that interactions returned by the stringApp API are undirected, these edges were introduced into the network in both directions.

Due to the implication of histone acetyltransferases (HATs) in the pathogenesis, a set of HATs was prioritized (data track “D”), and in conjunction with virus-implicated host proteins (VIHPs) it was utilized to construct a subnetwork with the help of SmartGraph. In the SmartGraph analysis, the HATs and VIHPs were used as starting and end nodes, and vice versa. In either case, the maximum length of shortest paths between starting and end nodes was limited to 3. For more details on the assembly of the SmartGraph subnetwork please refer to section *“Assembly of the SmartGraph Subnetwork”*, SI. This input dataset contains 248 HHIs involving 108 host targets (data track “I”).

The “Functional interactions” dataset (data track “K”), derived from Reactome database (version 2019) [6], [48] was used to cross-reference HHIs originating from various data sources.

### Drug–Target Interactions

Pertinent DTIs were primarily extracted from the DrugCentral database (version 2020, data track “E”) [10]. The DrugCentral database includes 4,642 drugs, of which 2,549 have regulatory approval dates, 3,082 have ATC codes [57], and 1,884 have INN stem annotations [58]. These drugs are associated with 110,577 formulations (drug labels) in total. Furthermore, the DrugCentral database also provides information on DTIs; 15,397 human DTIs and 4,910 nonhuman DTIs; of these 2,752 (2,328 human) are MoA drug-target associations.

For over 90% of the DTIs, the DTI therapeutic consequence is documented. We also collected the pharmacological action for each DTI which provides additional information about the potential intervention. The DTIs were originally extracted from scientific literature, drug labels and other data sources such as, ChEMBL [4], IUPHAR Guide2Pharmacology [59], WOMBAT-PK [60], DrugBank [61] and KEGG Drug [62].

To further enrich the network, we included both known and predicted DTIs for drugs and human endogenous metabolites found in the vicinities of the chemical space defined by three small molecules being investigated at the time for their antiviral activity against SARS-CoV-2 assays, namely, N4-hydroxycytidine (NHC) [63], hydroxychloroquine (HCQ) [64], [65] and camostat (CAM) [66]. A set of 25 compounds was compiled around NHC [67], 5 around the chemical neighborhood of HCQ, and camostat. Processing those 31 small molecules against the CLARITY platform [68] returned a total of 86 DTIs, known (from public sources [5]) and/or predicted [69] to have activity against 46 unique protein targets (data track “F”).

### Target Development Level Information

Target development level (TDL) information for host proteins were imported from the Pharos portal (data track “L”) [7], [70].

## Results and Discussion

### Semi-Automated Data Integration Workflow

In order to build a COVID-19 focused network, we needed to integrate data from multiple, diverse data sources. The bottleneck of the integration proved to be the consolidation of the differing data structures, exemplified by the lack of standardized data categories, preference of one protein identifier over the other, aggregating protein identifiers as delimiter separated list inside a column, to name a few. Also, some of the data originate from experiments whereas others from predictions. This made it necessary to prioritize data sources and to keep track of data provenance in a transparent manner. Here, we present a rigorous workflow that addresses the above challenges and can serve as a template for semi-automated integration of the future databases inspired by network pharmacology. Having such an integration workflow in place is key to the timely assembly of a network that is focused on a certain biological aspect, e.g. an infectious disease caused by an emerging pathogen.

The assembly of a COVID-19 focused network involved data sources that emerged over a matter of weeks since the start of the pandemic and others that were well-established long before. Considering that new information surfaced relatively quickly, we had to ensure the workflow we created was sufficiently flexible to accommodate new data. Here, we describe such a workflow (see: *Fig. 1*) and a COVID-19 focused Neo4j database that was produced by it. Information regarding the reproduction of the workflow is provided in the *“Reproducing the Integration Workflow*” section in SI.

The building blocks of the COVID-19 focused network represent experimentally determined as well as predicted HPI, HHI and DTI data, and additionally, prioritized host targets. In these data sources, host targets are typically identified by their gene name, with some exceptions where UniProt ACs are used, such as TDLs from Pharos DB and HHIs from SmartGraph. The name of several viral proteins in the two HPI datasets were slightly different. Such differences were manually reconciled (see: *“Mapping of Viral Protein Names in HPIs”* section in SI).

The next stage required harmonization of the input data structure. For each interaction type (HPI, HHI, DTI), a data structure was defined that was capable of accommodating all data of the respective interaction type. Naturally, each dataset needed to be tailored individually to fit the respective data structure, which is the main reason why the overall workflow is called semiautomated instead of automatic.

The origin of each data source was recorded and was assigned a priority value [71]. Of note, the choice of priority values is inherently subjective. However, these priority values can be easily reconfigured in the source code. As shown in *Fig. 1,* this step is the “workhorse” of the workflow, as it assures that practically any number of data sources can be integrated into a database, while preserving data provenance. The aim of annotating source priority is two-fold. First, it provides the means to deduplicate information in an unambiguous and reproducible manner. Second, it is possible to analyze only certain layers of the resultant database, by filtering data sources on the basis of their priority annotation.

The workflow of this study was designed to allow for the flexible data extension of data for each of the interaction types while providing an option to the investigator to restrict which data segment is subject to the extension. Briefly, *Fig. 1* depicts the following mechanism. Host proteins of potential importance were collected from data tracks “A”, “B”, “C”, “E”, “F”, “G”, and “H”. This set of unique proteins was used to induce an HHI subnetwork using the STRING and stringApp APIs [47], [56], [72] as described above. While an additional data source (data track “I”) contains HHIs extracted from SmartGraph, the implicated host proteins were not subject to the further STRING-expansion (unless there was an overlap between the proteins of these HHIs and the set of unique proteins extracted above). Once the induced HHI subnetwork was returned by the STRING API, it was also subject to data structure harmonization and data provenance annotation (data track “J”).

Merging the different types of interactions, namely HHIs, HPIs and DTIs, was easy due to the central harmonization mechanism. Data deduplication was carried out with the help of priority annotations. In particular, multiple occurrences of an interaction between identical entities were aggregated to keep only one interaction annotated with the highest priority data source. Of note, each source of the interaction in question was recorded as metadata, in order to maintain data provenance. This metadata provides an additional layer of control to analyze the integrated database.

One of the final challenges that we had to address in the workflow was integration of HHIs from different sources. Typically, this procedure is rather challenging due to potentially conflicting and missing HHI information. For instance, HHIs from the STRING API are not annotated with the mode of regulation (whether a target up-regulates or down-regulates its interacting partner).

A viable strategy to resolve such issues is to utilize a comprehensive HHI database as an external reference. Although the aforementioned step of HHI deduplication based on the data source priority assures that HHI information of higher reliability is retained, using an external HHI data source can add more confidence to data reconciliation – more so, if the external data source is curated. We decided to use the “functional interactions” subset of Reactome (RFI) to fulfill this role. Each HHI was cross-referenced to the RFI subset. This process resulted in the assignment of both the direction and mechanism of regulation for each HHI in the integrated database, as well as the respective confidence score for each RFI. If a given HHI was missing from the RFI subset or the mechanism or the direction of regulation was undefined, then the corresponding properties of that HHI were set to “unknown”. Of note, metadata extracted from the original data source was preserved for all HHIs to maintain data provenance.

Lastly, each protein target was annotated by the respective TDL category extracted from the Pharos [7] database. However, as proteins are encoded with UniProt ACs [73], [74] both in Pharos and in SmartGraph, we had to resolve them to gene names and *vice versa.* We used the UniProt API [75] to retrieve UniProt to gene name mapping. Considering that this mapping is a many-to-many relationship, the 1:1 mapping was achieved by retaining the highest TDL of any of the UniProt ACs in the case of Pharos data. The TDL annotation of targets enables investigators to instantly identify targets for which FDA-approved drugs exist. This information combined with pathway analysis provide the foundation for the formulation of drug repositioning hypotheses. Such hypotheses can mean the first steps towards the discovery of therapeutics.

## Database Deployment and Dissemination

### Neo4COVID19 Database

The data integration scheme discussed above was implemented in Python [76] and is available as a source code repository at https://github.com/ncats/neo4covid19 [77]. The data integration script builds a Neo4j database [78] which can be accessed publicly via Neo4j Browser, a web-based graphical user interface (GUI) provided by Neo4j or using various Neo4j API interfaces, e.g. “py2neo” [79], which make use of the Neo4j Bolt protocol [78]. Details on how to access the database are available on the Neo4COVID19 website at https://neo4covid19.ncats.io.

As Neo4COVID19 is a graph-database, fundamentally, it is a collection of nodes and edges. In the database, there are 950 “Target” nodes and 641 “Compound” nodes representing proteins and drugs, respectively. With the help of node attributes, Target nodes are further differentiated into 55 viral and 895 host Targets. Two types of relationships are defined between nodes: “Interacts” and “DTI”. Each relationship of the former type involves two host proteins (74,805 HHIs) or a host and a viral protein (487 HPIs). DTI relationships (1,265) are defined between Compound and Target nodes. Further information regarding the node and edge types extracted from each data resource are provided in *Table 1.*

**Table 1.** COVID-19 focused network statistics. Shown are summary of individual data types integrated into the Neo4COVID-19 Neo4j database. Of note, overlap may exist between data types associated with the original data sources. HHIs: host–host protein interactions, HPIs: host– pathogen (here: SARS-CoV-2) protein interactions, DTIs: drug–target interactions.

| Dataset | Host Targets | Viral Targets | Compounds | HPIs | HHIs | DTIs |
| --- | --- | --- | --- | --- | --- | --- |
| Proteomics Study | 102 | - | - | - | - | - |
| CRISPR | 105 | - | - | - | - | - |
| Meta Path AI/ML | 185 | - | - | - | - | - |
| STRING | 793 | - | - | - | 74,584 | - |
| SmartGraph / HATs | 116 | - | - | - | 241 | - |
| Interactome Study | 332 | 27 | - | 332 | - | - |
| P-HIPster | 38 | 28 | - | 155 | - | - |
| Predicted DTIs | 46 | - | 31 | - | - | 86 |
| DrugCentral | 129 | - | 625 | - | - | 1,207 |

Node and edge attributes shown in *Table S1-S2* in SI can be used to fine-tune the network in a way that matches the needs of the analysis at hand. For instance, one might decide to only consider nodes and edges of the COVID-19 focused network that represent experimental data. This can be achieved with ease by filtering the data with the help of Boolean fields, each representing a specific data source. Another example is application of a confidence threshold to HHI edges retrieved from stringApp API using the “source_specific_score” field value to filter the data. This data structure facilitates the versatile use of the Neo4COVID19 database in many research settings and the corroboration of information pertaining to various interactions.

### Importing Neo4COVID-19 Database into Cytoscape

In order to facilitate the translational impact via dissemination [80] and flexible downstream analysis of the Neo4COVID19 focused network in the bioinformatics community, here we describe a simple procedure to import the database into the widely utilized Cytoscape application [81].

Importing the Neo4COVID19 database into Cytoscape requires the installation of the “Cytoscape plugin for Neo4j” [82] which can be easily achieved from within the Cytoscape application (see: *Fig S1a* in SI). Once the plugin is installed, a connection to the Neo4j database has to be established via the Bolt protocol, as shown in *Fig S1b* in SI.

After that, the successful establishment of database connection, the entire Neo4COVID19 database can be imported into Cytoscape with only a basic query statement written in the Cypher [78] language as shown in *Fig S1c* in SI. The statement is actually identical to the query provided by the plugin when removing the “LIMIT” clause from the default statement. Finally, a custom visualization style can be applied (see: *Fig S2* and “Applying Custom Visual Style to the Imported Network in Cytoscape”, SI). The resultant network is shown in *Fig S1d* in SI.

## Use Cases

Here, we describe use cases to demonstrate how one can use the Neo4COVID19 for hypothesis generation in a network pharmacology setting.

First, we examined the T_clin_-designated HPI subset: T_clin_ designated proteins (according to TDL) from the Krogan dataset [ref] that have a fold change of 10 or higher following SARS-CoV-2 exposure. Out of 166 such HPIs, at least 63 occur between 27 human and 11 viral proteins. These 27 proteins are integral components of the mitochondrial respiratory chain complex I (GO:0005747), and are targeted by the antidiabetic drug metformin. Initially, we were excited to note that metformin shares chemical similarity with an old antiviral drug moroxydine (see: *Fig 3*). Shortly thereafter, as type 2 diabetes was identified as risk factor for severe COVID-19 [83], we had to briefly suspend further metformin research as we suspected its effect on the T_clin_-designated HPI subset was indirect. However, recent reports show that metformin treatment is actually independently associated with a significant reduction in mortality in subjects with diabetes and COVID-19 [84]. Thus, while we cannot assume that the antiviral activity of moroxydine is similar, we are quite confident that metformin acts as a significant HPI perturbagen during SARS-CoV-2 infection and could serve as an adjuvant antiviral therapy under appropriate conditions.

**Figure 2.**
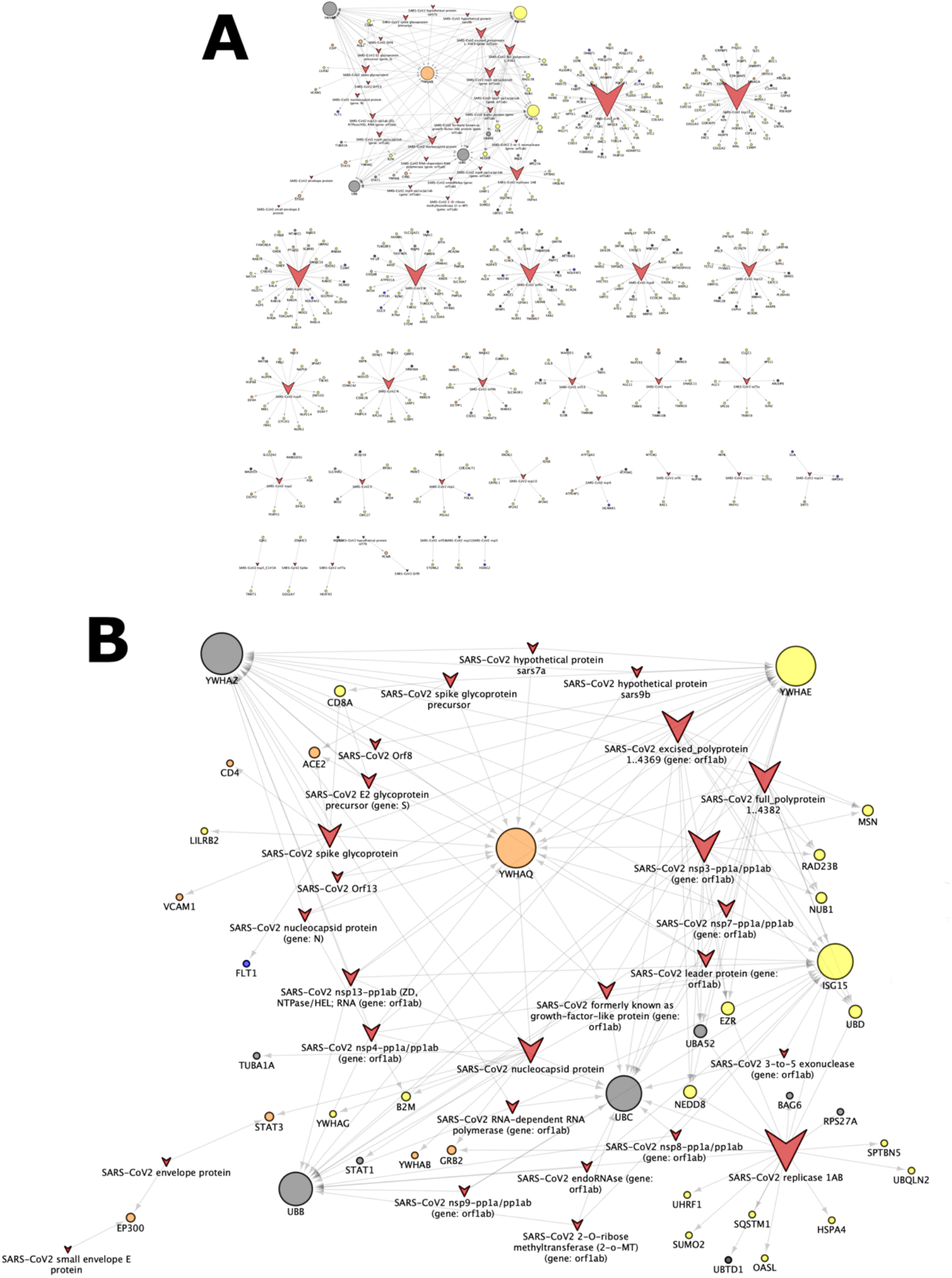
Bipartite network of HPIs. Human and virus proteins are depicted by circles and “v-like” shapes, respectively. The larger the node size, the higher the degree of the node connectivity. Color of the human proteins encode their TDL annotation: blue: T_clin_, orange: T_chem_, yellow: Tbio, dark gray: Tdark, light gray: unknown. **A:** The complete HPI bipartite network. **B:** The subnetwork centered around the virus hub YWHAQ. The network was visualized with the help of Cytoscape v3.6.0 [86].

**Figure 3.**
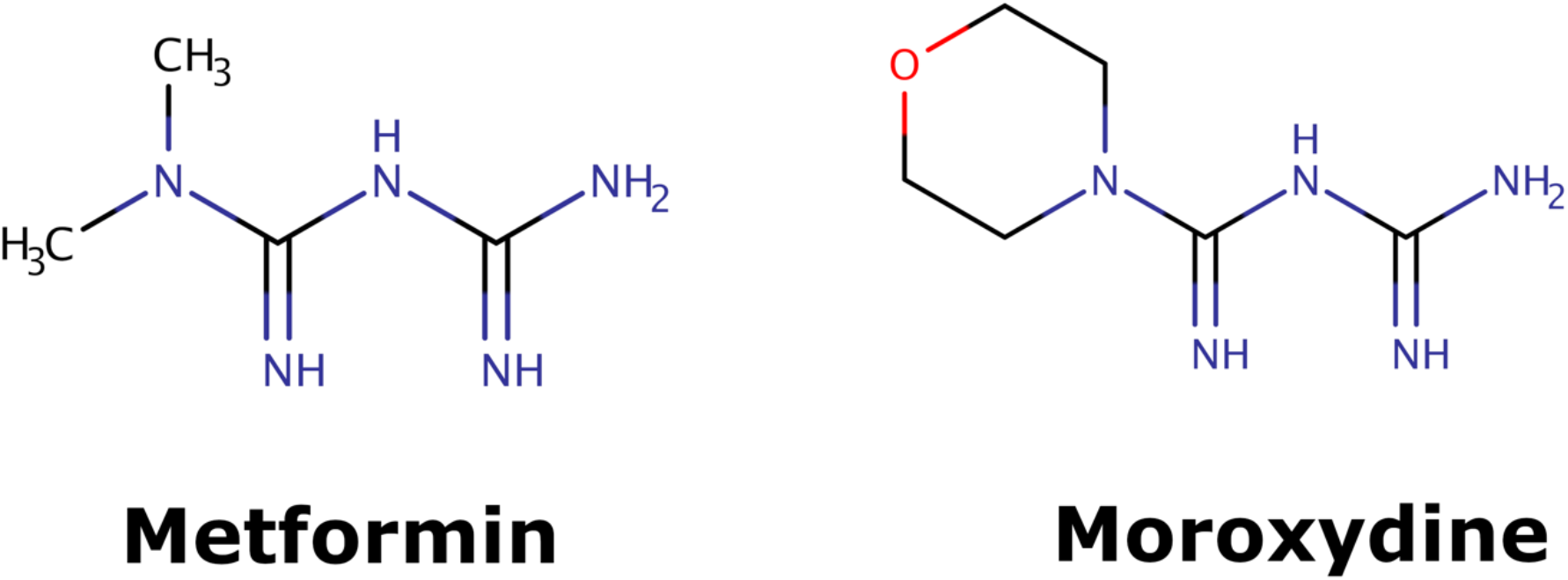
Molecular structures of metformin and moroxydin. Molecules were depicted with the help of ChemAxon’s MarvinSketch v17.15.0 [87].

Next, we investigated the interactions between host and virus proteins (the HPI relationships). With the help of node and edge attributes it is possible to construct a sub-network of the Neo4COVID19 network by retaining only virus and host targets, and the edges between them. This process gives rise to a bipartite network, in which host nodes are only connected to virus nodes and *vice versa*. In this network, a natural clustering emerges where several connected components exist involving many human proteins centered around a single virus protein (see: *Fig 2a*.). The network topology also reveals that certain host targets might be thought of as the *“Achilles’ heel”* of the virus due to their connection structure. The peculiarity of these host proteins is that they are connected to many virus proteins hence they were named as “virus hubs”. An ideal strategy would be to target such virus hub that affects multiple biological processes of the virus while only causing a small perturbation in the regulatory network of the host.

As shown on *Fig 2b,* a few host targets are connected to a substantially higher number of nodes than others, as indicated a larger size of the corresponding nodes. One of such targets, YWHAQ (UniProt AC: P27348), is associated with “T_chem_” TDL category, which means that there is a small molecule modulator for that target. Therefore, one might hypothesize that YWHAQ might be a potential host target to develop a drug against based on the existing modulator(s) as seed active molecule(s). However, the next step will require the analysis of the role of YWHAQ in the signal transduction network of the host (human). While such analysis is outside the scope of this work, it should be noted that the Pharos database indicates 260 HHIs associated with YWHAQ, which suggests that this particular target is involved in many biological processes of the host, hence its modulation will likely considerably perturb the network. It could be still possible that a redundant (parallel) paths exist in the signal transduction network, which might mitigate the potential adverse effects of the perturbation.

While the aforementioned strategy might identify promising virus hubs, unfortunately, there are no T_clin_ targets among them. This indicates that at the time of this study we could not apply drug repositioning to target virus hubs. An alternative strategy could be to target multiple T_clin_ host targets (blue circles on *Fig 2a*) that appear in separate clusters. Drug compounds associated with T_clin_ targets can be easily imported into the network. This approach might form the basis for the engineering of a multi-agent therapy ideally employing approved drugs if they can effectively interfere with the implicated pathogen-host interactions.

The detailed procedure to replicate the uses cases described above is provided in “Reproducing the Use Cases” section in SI.

## Considerations Regarding the Validity of the Data

The data integration workflow presented in this study was designed to be widely adoptable in a network pharmacology research setting. In this sense, our workflow can facilitate any semiautomated integration of constantly evolving data – which happens to be the nature of many modern data sets, such as Reactome [6], DrugCentral [10], and Pharos [7], to name a few. However, the unprecedented pace of data influx we witnessed at the beginning of the COVID-19 pandemic highlighted a pitfall that is more related more to data quality than to the complexity of data integration.

Specifically, the validity of some early COVID-19 related results have since been questioned, and manuscripts have been retracted [65], [85]. We contemplate that the dichotomy between the need for relevant and bleeding edge information and the quality of the data will recreate this scenario in the case of a similar event in the future. Therefore, any data processing workflow should be dynamic, transparent, regularly reviewed, and constantly updated.

## Conclusions

Here, we describe a semi-automated workflow for the integration of data sources to produce a COVID-19 focused graph database. The workflow can be easily generalized to other drug discovery scenarios which can save precious time in the case of a pathogen outbreak. The workflow makes use of the state-of-the-art network pharmacology approaches and yields an interconnected network of host and viral protein targets and drugs, containing information on HPIs, HHIs, and DTIs. The workflow is flexible, which makes it possible to replace data sources and/or add new ones to it. The data aggregation process can also be altered by redefining the priority of individual data sources. Furthermore, the layered structure of the network and the underlying data schema allows researchers to filter data sources that they find relevant in their investigation.

During the development of this workflow we came across known bottlenecks related to data integration that we believe could be ameliorated to a great extent by following certain practices. For instance, the interaction types in a network pharmacology setting are well defined (HHI, HPI, DTI), and therefore, such data should be made available in a few well-established formats. This would allow for seamless integration of already existing and emerging datasets. Furthermore, providing a robust API for a dataset facilitates its integration and allows for programmatic updates.

The Neo4j database generated by this workflow can be accessed via a web interface at https://neo4covid19.ncats.io to enable the exploration of data without much expertise in the bioinformatics field. In addition, it takes advantage of the Neo4j Bolt protocol and provides an API to facilitate the integration of the COVID-19 focused network into virtually any bioinformatics workflow. We provided use cases to show how the Neo4COVID-19 network can be utilized to generate hypotheses with focus on drug repositioning.

We believe that our Neo4COVID19 database will be a valuable asset to the research community and will catalyze the discovery of therapeutics to defeat COVID-19. Furthermore, the underlying flexible workflow can serve as a starting point for the integration of critical knowledge in the event of a potential future outbreak, which we all hope will never happen.

## Abbreviations

**Abbreviations not defined in the text are listed below**
API: Application Programming Interface
ATC: Anatomical Therapeutic Chemical
INN: International Nonproprietary Names
MoA: Mechnism-of-Action
FDA: U.S. Food and Drug Administration

## Declarations

### Contributions of Authors

This research study was initiated by TIO and GZK. The workflow and the Neo4j database was designed and built by GZK. MG, BG and BM designed and configured the computational infrastructure to provide public access to the Neo4j database. AGG, SGM, EM, DM and MDH provided inspiration and feedback for the study. TIO, PK, JM and LJJ provided predictions for HPIs, host and viral targets, and drugs. TIO, PK, LJJ and VBS contributed with data analysis. GZK and VBS wrote the majority of the text, TIO, LJJ, JM, VBS, PK, MG, IG, LB, MDH, AGG provided edits to the manuscript, the others contributed to the study. All authors read and approved the manuscript.

## Acknowledgements

The authors are thankful for the devoted work of Henderson Tozer, Kevin Diaz, James McWilliams and all members of the IT support team of the Information Resources Technology Branch of NCATS/NIH who enabled us to continue our research in unprecedented times. Furthermore, we would like to acknowledge Tim Willson, Dac-Trung Nguyen and Noel T. Southall, PhD for their fruitful discussions.

## Availability of Data and Materials

The Neo4COVID19 database is publicly available at https://neo4covid19.ncats.io where information is also provided on how the database can be accessed via the BOLT protocol using API.

The source code repository of the workflow utilized to construct the Neo4COVID19 database is publicly available at https://github.com/ncats/neo4covid19.

