## Supporting Information for "A Workflow of Integrated Resources to Catalyze Network Pharmacology Driven COVID-19 Research"

### Sample Python Code Snippet to Access Neo4COVID19 Database via API

Details on how to install the "py2neo" Python library [1], [2] are provided at https://py2neo.org/v4/.

Sample Python code snippet to connect to the Neo4j database and retrieve the result of the CYPHER query [3].

```
from py2neo import *
graph = Graph(host="neo4covid19.ncats.io", bolt_port=7687, user='', password = '', secure = True)
graph.run("MATCH (t:Target) RETURN t LIMIT 5").data()
```

Furthermore, a script distributed as part of the https://github.com/ncats/neo4covid19 source code repository [4] provides specific examples to query the Neo4COVID19 database. The file is located under `neo4covid19/code/generate_stats.py` where the "`neo4covid19/`" part of the path is the root of the repository.

### Pseudo-Code of the Data Integration Workflow

Here we provide the pseudo-code of the data integration workflow conceptualized by *Fig 1*. The name of the variables associated with input data sources is identical to the label of the respective data track.

*Algorithm*

```
Input: data frame A                    // HPIs
Input: data frame B                    // HPIs
Input: data frame C                    // host proteins
Input: data frame D                    // host proteins
Input: data frame E                    // DTIs
Input: data frame F                    // DTIs
Input: data frame G                    // host proteins
Input: data frame H                    // host proteins
Input: data frame K                    // HHIs
Input: data frame L                    // TDLs


Variable: map (String, Int) priorityMap{}
Variable: data frame allHPIs
Variable: data frame allHHIs
Variable: data frame allDTIs
Variable: data frame uniqueHostProteins
Variable: data frame uniqueVirusProteins
Variable: data frame uniqueDrugs
Variable: data frame proteins
Variable: data frame I_forward         // HHIs from SmartGraph (forward direction)
Variable: data frame I_reverse         // HHIs from SmartGraph (reverse direction)
Variable: data frame I                 // HHIs from SmartGraph
Variable: data frame J                 // HHIs from STRING


priorityMap = assignPriorityValues ([A, B, C, D, E, F, G, H, I, J])

[A, B] = harmonizeVirusProteinIdentifiers ([A, B])

[A, B] = harmonizeHPIDataStructure ([A, B])

[C, D, G, H] = harmonizeHostProteinDataStructure ([C, D, G, H])
```

[*E, F*] = harmonizeDTIDataStructure ([*E, F*])

[*A, B, C, D, E, F, G, H*] = recordDataProvenance ([*A, B, C, D, E, F, G, H*])

[*A, B, C, D, E, F, G, H*] = annotateDataSourcePriority (*priorityMap,* [*A, B, C, D, E, F, G, H*])

*allHPIs* = appendByRows([*A, B*])

*allHPIs* = deduplicateByPriority (*allHPIs*)

*uniqueHostProteins* = extractUniqueHostProteins ([*allHPIs, C, F, G, H*])

*uniqueVirusProteins* = extractUniqueVirusProteins ([*allHPIs*])

*allDTIs* = appendByRow ([*E, F*])

*allDTIs* = deduplicateByPriority (*allDTIs*)

*uniqueDrugs* = extractUniqueDrugs (*allDTIs*)

*J* = extendHHIsByStringApp (*uniqueHostProteins, species_ncbi = 9606, limit_of_mapped_genes = 1, max_interactor = 100, score_cutoff = 0, alpha = 0.5*)

*I_forward* = expandHHIsBySmartGraph (*D*, *uniqueHostProteins, maxDistance = 3, minConfidence = 0*)

*I_reverse* = expandHHIsBySmartGraph (*uniqueHostProteins, D, maxDistance = 3, minConfidence = 0*)

*I* = appendByRow (*I_forward, I_reverse*)

[*I, J, K*] = harmonizeHHIDataStructure ([*I, J, K*])

[*I, J, K*] = recordDataProvenance ([*I, J, K*])

[*I, J*] = annotateDataSourcePriority (*priorityMap, I, J*)

*allHHIs* = appendByRow ([*I, J*])

*allHHIs* = deduplicateByPriority (*allHHIs*)

*allHHIs* = overlayReferenceHHIData (*allHHIs, K*)

*proteins* = extractUniqueHostProteins ([*I, J*])

*uniqueHostProteins* = appendByRows([*uniqueHostProteins, proteins*])

*uniqueHostProteins* = appendByRows([*uniqueHostProteins, proteins*])

*uniqueHostProteins = annotateTDL (uniqueHostProteins, L)*

populateNeo4jDatabase (*uniqueHostProteins, uniqueVirusProteins,*
*uniqueDrugs, allHHIs, allHPIs, allDTIs*)

### Reproducing the Integration Workflow

In order to reproduce the workflow, provided the required Python [1] environment has been set up, a local copy of the `neo4covid19` repository needs to be created as follows.

```
git clone https://github.com/ncats/neo4covid19
```

Note, that paths referring to files in this manuscript start with "`neo4covid19`". In this context, `neo4covid19` points to the root directory of the local copy of the cloned repository.

The first stage of the workflow is executed as:

```
python prepare.py
```

In case an error occurs due to an API call, try this command instead:

```
python prepare.py test
```

This is followed by assembling the SmartGraph subnetwork. For details, please refer to section "*Assembly of the SmartGraph Subnetwork*".

The last stage of the workflow is executed as:

```
python compile.py
```

Or, in the case of an API call error:

```
python compile.py test
```

### Assembly of the SmartGraph Subnetwork

In order to reveal potential connection between histone acetyltransferases (HATs) and SARS-CoV-2 virus implicated host proteins (VIHPs), we performed network analysis with the help of the SmartGraph platform [5]. Since a set of VIHPs is compiled in the integration workflow, it was necessary to implement a breakpoint in the workflow. Upon completion of the first part of the workflow, SmartGraph analysis is performed, and the results are subsequently fed to the second stage of the workflow to finish the integration. While this scenario is not ideal, at the time of the workflow creation, the SmartGraph platform did not provide API access.

The gene names of VIHPs were mapped to UniProt IDs [6], [7] to comply with the SmartGraph input requirements. First, VIHPs present in the file `chembl_uniprot_mapping.txt` (distributed as part of ChEMBL database, version 27 [8]) were identified. Next, with the help of UniProt (API) [7], [9] the UniProt IDs of these genes were retrieved.

These are the detailed step to assemble the SmartGraph subnetwork. Assuming you have created a local copy of the neo4covid19 repository (see above), perform the following steps:

1. Go to SmartGraph (https://smartgraph.ncats.io).
2. Clear the fields "Start Nodes" and "End Nodes" then click on "clear graph".
3. Copy the IDs in column ‘uniprot_id’ of file `neo4covid19/data/input/HATs.tsv` (note that the “`neo4covid19`” points to the root of the `neo4covid19` repository). Insert this set of UniProt IDs as "Start Nodes" in SmartGraph (https://smartgraph.ncats.io).

4. Copy the UniProt IDs from the output of Step 1 located at `neo4covid19/data/output/unique_host_proteins_prestring.txt`. Copy the UniProt IDs and insert them as "End Nodes" in SmartGraph.
5. Set the "Max Distance" parameter to 3.
6. Leave the "PPI Confidence Level" to its default value, i.e. 0.00.
7. Click on "find shortest path".
8. Once the network is assembled in SmartGraph, click on "Download graph", select "Cytoscape JSON", then rename the downloaded file to `SG_HATs_dist_3_conf_0.00.json` and place the file into `neo4covid19/data/input/`.
9. Repeat steps 2-7 but this time use the HATs as “End Nodes” and the UniProt IDs in `neo4covid19/data/output/unique_host_proteins_prestring.txt` as “Start Nodes”.
10. Save the resultant network in "Cytoscape JSON" format and save it as `SG_HATs_reverse_dist_3_conf_0.00.json` and place the file into `neo4covid19/data/input/`.

### Expansion of HHIs via StringApp API

Expanding the HHIs present in a preliminary Neo4COVID-19 network was performed in a two-step procedure employing the STRING [10] and stringApp APIs [11].

In the first step, the gene symbols of human proteins in pre-expanded Neo4COVID-19 network were translated into the STRING database identifiers with the STRING API. We utilized the following URL for this API call: https://string-db.org/api/tsv-no-header/get_string_ids . Gene symbols were passed to parameter `identifiers` as a newline "\n" separated string (without quotation marks). Mapping of gene identifiers was forced to a one-to-one mapping by selecting the "best" STRING ID for a given gene symbol by setting `limit` to 1. In addition, we limited the mapping to human genes only by setting `species` to 9606; we included the original IDs in the results by setting `echo_query` to 1; and we provided a string to our liking for `caller_identity`.

Next, with the returned STRING database IDs we made a second API call to URL `https://api.jensenlab.org/network` . The STRING database IDs were passed to the `entities` parameter as a newline "`\n`" separated string. The `additional` parameter was set to 100, which defines the maximal number of proteins the original network can be extended with. Parameter `alpha` was set to its default value of 0.5.

The basis of the expansion is the computation of a connectivity score for proteins not in the query network. The connectivity score is a ratio of the total connectivity score of a given protein to the query proteins versus its total connectivity score to all proteins in STRING database [Ref].

For more details, please refer to the section “Network Expansion” in the study of Doncheva *et al*. [11].

Of note, the following genes present in the pre-extension network were excluded from the STRING extension process as they produced errors when included into the API call: ELOC, EP300, SLC25A5, TUBA1A, STAT1, ELOB, RBX1, CREBBP, SKP1.

#### Applying Custom Visual Style to the Imported Network in Cytoscape

The file containing the custom Cytoscape [12] visual style (style_Neo4COVID19.xml) is distributed as part of the Neo4COVID19 code repository (`neo4covid19/code/style_Neo4COVID19.xml`) [4]. The process of importing and applying the custom style is shown on Fig S2.

##### Mapping of Viral Gene Names

We have established a mapping between the viral gene names predicted by P-HIPSTer [13], [14] and those reported in the interactome study by [15], [16] The mapping is provided on sheets “ID_Mapping” and “Sheet1_MappedIDs” in the file `data/output/Merged.xlsx` in the `neo4covid19` repository [4].

### Reproducing the Use Cases

1. Network assembly

   - Establish network connection:

     `Apps > Cypher Queries > Connect to Neo4j Instance`

     Provide `aspire.covid19.ncats.io:7687` as `Hostname`, leave rest of the form empty, then click on `Connect`.

   - Import bipartite HPI network

     `Apps > Cypher Queries > Import Cypher Query`

     Enter this Cypher Query:

     ```
     match (n)-[r:INTERACTS]->(m) WHERE r.interaction_type="HPI" return n,r,m
     ```

     Click on `Execute Query`.

2. Apply visual style

   - Please refer to *"Applying Custom Visual Style to the Imported Network in Cytoscape"* section in SI.

3. Topology analysis

   - `Tools -> NetworkAnalyzer -> Network Analysis -> Analyze Network ...`

     Select `Treat the network as directed.`, click on `OK`

4. Adjust node size as a function of "EdgeCount"

   - Click on `Style` on the left panel and select `Neo4COVID19` in the drop-down box.
   - Click on `Node` on the bottom of the visualization panel.
   - Select `Size`, set `Column` to `EdgeCount`, then set `Mapping` to `Continuous Mapping`.
   - Adjust the gradient as shown on the small panel until there is a good separation between low and high degree nodes.

### Figures

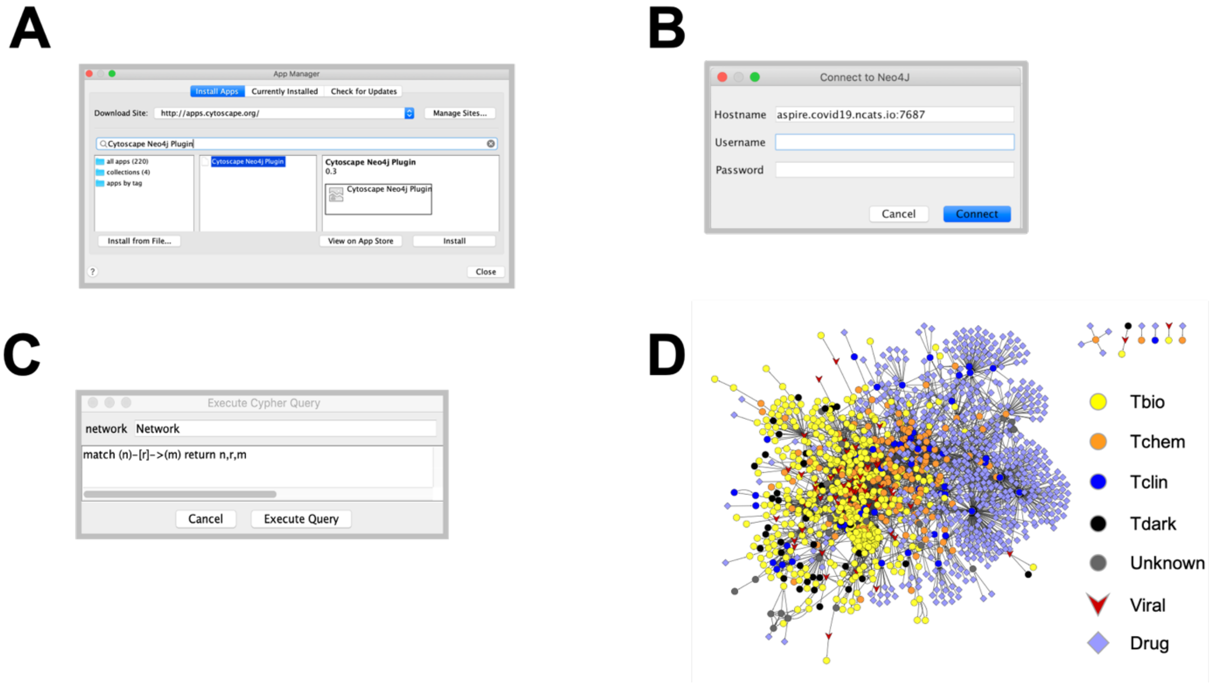


**Figure S1. Process of importing the COVID-19 focused network from Neo4j into Cytoscape. A)** Installing the "Cytoscape Neo4j Plugin" [17] by navigating to `"Apps -> App Manager…"`, typing "Cytoscape Neo4j Plugin" in the search bar, selecting the plugin from the results and finally clicking `"Install"`. **B)** Establishing Neo4j database connection ("`Apps > Cypher Queries > Connect to Neo4j Instance`"). Note, that neither username nor password is required. Host: `aspire.covid19.ncats.io:7687` . **C)** Cypher query to import the entire Neo4COVID19 network into Cytoscape ("`Apps > Cypher Queries > Import Cypher Query`", query: `match (n)-[r]->(m) return n,r,m` ) . **D)** Resultant network (after applying the custom visual settings). Nodes representing host and viral proteins, and drugs are denoted by circle, "V", and diamond shaped nodes. Where applicable, the target development category (TDL) [18], [19] of proteins are color-coded according to legend. Screenshots were made from the Cytoscape application.

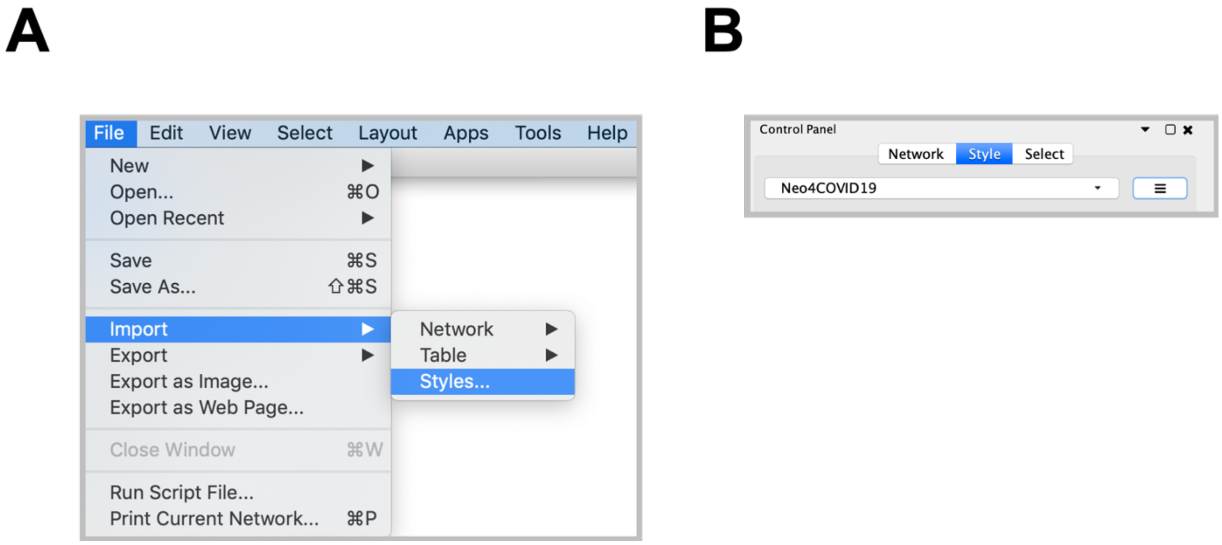


**Figure S2. Customizing network visualization. A)** Importing the "style_Neo4COVID19.xml" file that contains the custom visual style definition. **B)** Applying the custom visual style "Neo4COVID19".

### Tables

| Target | | Compound | |
|---|---|---|---|
| *attribute* | *type* | *attribute* | *type* |
| gene_symbol | string | drug_name | string |
| target_type | string | smiles | string |
| tdl | string | inchi | string |
| uniprot | string | inchi_key | string |
| is_in_preprint | boolean | ns_inchi_key | string |
| is_in_phipster | boolean | CAS_RN | string |
| is_in_taiml | boolean | struct_id | string |
| is_in_jdti | boolean | is_in_drugcentral | boolean |
| is_in_string | boolean | is_in_jdti | boolean |
| is_in_hats | boolean | is_in_hcq | boolean |
| is_in_natdt | boolean | is_in_nhc | boolean |
| is_in_drugcentral | boolean | is_in_cam | boolean |
| is_in_crispr | boolean | is_in_drugs | boolean |
| metadata | string | metadata | string |
| uuid | string | uuid | string |

**Table S1. Node attributes of the Neo4COVID19 graph database.**

| Interacts | | DTI | |
|---|---|---|---|
| *attribute* | *type* | *attribute* | *type* |
| source_node | string | edge_label | string |
| target_node | string | drug_name | string |
| interaction_type | string | source_node | string |
| interaction | string | target_node | string |
| mechanism | string | action_type | string |
| reactome_mechanism | string | p_chembl | numeric |
| reactome_regdir | string | is_activity_known | boolean |
| reactome_score | numeric | priority | integer |
| metadata | string | source | string |
| comment | string | relationship | string |
| pmids | string | comment | string |
| priority | integer | pmids | string |
| source | string | metadata | string |
| data_origin | string | is_in_drugcentral | boolean |
| relationship | string | is_in_jdti | boolean |
| source_specific_score | numeric | source_node_uuid | string |
| is_in_preprint | boolean | target_node_uuid | string |
| is_in_phipster | boolean | uuid | string |
| is_in_string | boolean | | |
| is_in_hats | boolean | | |
| is_in_reactome | boolean | | |
| source_node_uuid | string | | |
| target_node_uuid | string | | |
| uuid | string | | |

**Table S2. Edge attributes of the Neo4COVID19 graph database.**

### References


[1] “Python Core Team. Python: A dynamic, open source programming language. Python Software Foundation.” https://www.python.org/.

[2] “Python Library ‘py2neo’ v4.” .

[3] “Neo4j Graph Database.” https://neo4j.com/.

[4] “Code Repository ‘neo4covid19.’” https://github.com/ncats/neo4covid19.git.

10.1093/nar/gky1131.

[11] N. T. Doncheva, J. H. Morris, J. Gorodkin, and L. J. Jensen, “Cytoscape StringApp: Network Analysis and Visualization of Proteomics Data,” *J. Proteome Res.*, vol. 18, no. 2, pp. 623–632, Feb. 2019, doi: 10.1021/acs.jproteome.8b00702.

[12] P. Shannon *et al.*, “Cytoscape: a software environment for integrated models of biomolecular interaction networks.,” *Genome Res.*, vol. 13, no. 11, pp. 2498–504, Nov. 2003, doi: 10.1101/gr.1239303.

[13] G. Lasso *et al.*, “A Structure-Informed Atlas of Human-Virus Interactions,” *Cell*, vol. 178, no. 6, pp. 1526-1541.e16, Sep. 2019, doi: 10.1016/j.cell.2019.08.005.

[14] “P-HIPSTer.” http://phipster.org/.
